# Predicting Collapse of Complex Ecological Systems: Quantifying the Stability-Complexity Continuum

**DOI:** 10.1101/713578

**Authors:** Susanne Pettersson, Van M. Savage, Martin Nilsson Jacobi

**Affiliations:** Department of Space, Earth and Environment, Chalmers University of Technology, Maskingränd 2, 412 58 Gothenburg, Sweden; Department of Ecology and Evolutionary Biology, Department of Biomathematics, UCLA, Los Angeles, CA, 90095, USA

## Abstract

Dynamical shifts between the extremes of stability and collapse are hallmarks of ecological systems. These shifts are limited by and change with biodiversity, complexity, and the topology and hierarchy of interactions. Most ecological research has focused on identifying conditions for a system to shift from stability to any degree of instability—species abundances do not return to exact same values after perturbation. Real ecosystems likely have a continuum of shifting between stability and collapse that depends on the specifics of how the interactions are structured, as well as the type and degree of disturbance due to environmental change. Here we map boundaries for the extremes of strict stability and collapse. In between these boundaries, we find an intermediate regime that consists of single-species extinctions, which we call the Extinction Continuum. We also develop a metric that locates the position of the system within the Extinction Continuum—thus quantifying proximity to stability or collapse—in terms of ecologically measurable quantities such as growth rates and interaction strengths. Furthermore, we provide analytical and numerical techniques for estimating our new metric. We show that our metric does an excellent job of capturing the system behaviour in comparison with other existing methods—such as May’s stability criteria or critical slowdown. Our metric should thus enable deeper insights about how to classify real systems in terms of their overall dynamics and their limits of stability and collapse.

---

SYSTEM stability and collapse are core concepts for ecology and complex systems that have been studied both theoretically and empirically. The emerging picture of stability is multi-faceted. Over the years many features of ecosystems have been posited as stabilising factors such as restricted number of trophic levels [1], hierarchical interactions [2], compartmentalisation [3], existence of specific species interaction motifs among the interactions [4, 5], long weakly-interacting trophic loops [6], large numbers of prey or predators per species [7], number of mutualistic partners [8], structural asymmetry [9], nestedness [10], species body-size ratios [11], species functional complementarity [12], correlations in species interaction strengths [13], trophic coherence [14], and adaptive foraging [15]. Moreover, some features are not purely stabilising or destabilising and can interact in non-trivial ways [16, 17]. In addition, there are multiple perspectives on stability, including resilience and resistance [18, 19] as well as many ways to represent the web of interactions between species in an ecosystem (trophic [9, 11, 20], mutualistic [21, 22], antagonistic [23], competitive [24], mixed [25, 26] and multilayer [27]).

Properties of ecosystem stability can be evaluated using dynamical models. Two examples are the analysis of stabilising effects of interaction modules with few species [1, 28], and modelling systems with large numbers of species [29–31]. Pioneering work on system stability was done by Robert May who used Random Matrix Theory to show the importance of system complexity measured by biodiversity (defined as species richness), number of interactions, and variance of interaction strengths [32]. In May’s model complexity can be detrimental for stability, an insight that ran counter to the previous paradigm that complexity begets stability through functional redundancies or because of less reliance on a single keystone species [33, 34]. This paradigm was based on observations that ecosystems with few species, like agricultural soils, could collapse due to large fluctuations in population abundances of pests. In contrast, species-rich and highly complex ecosystems, like the Amazon rain-forests, had not been observed to exhibit such large fluctuations [35].

One of May’s key contributions was to introduce a complexity boundary where an ecosystem loses stability. The existence and location of a complexity boundary has been the focus of nearly five decades of work and is still an ongoing debate. Examples of recent studies include biologically-inspired interactions, such as modular [16], exclusively predator-prey interactions [36] and interaction diversification [31]. Some of these features shift the boundary to higher levels of complexity, but an upper limit still exists above which the system will lose stability.

There are several different criteria for stability that are related to the equilibrium state of species abundances. In the context of this paper, resilience means that the system will return to the exact same equilibrium state following a perturbation in species abundances. Another type is structural stability, which means that small changes in system properties – growth rates, interaction structure – do not lead to drastic changes, such as species extinctions [37]. Persistence is another aspect of stability, measured by the fraction of species out of the initial biodiversity that are present at equilibrium [3, 21, 38]. These criteria have been discussed extensively in the literature but rarely in concert [39, 40].

Here, we revisit random interaction matrices to develop a framework that includes all these aspects of stability. We show that taking persistence into consideration in the analysis of complexity boundaries typically leads to two boundaries, instead of one. The first boundary is between Strict Stability (SS), where systems are both structurally stable and resilient, and an Extinction Continuum (EC) where systems are structurally unstable but still resilient. The second boundary is between the Extinction Continuum and Collapse (C) with possible chaotic dynamics, limit cycles, or a new fixed point with substantial loss of species. As such, the Extinction Continuum represents an intermediate, previously overlooked but ecologically important regime between these two boundaries.

Furthermore, we can estimate the location of the two boundaries, the system’s persistence, and proximity to collapse in the Extinction Continuum, based only on measurable quantities of real systems. In contrast to previous studies we do not rely on the initial biodiversity that cannot be observed [40, 41]. Thus, we extend the current knowledge and toolkit by providing a framework and a metric that can predict a system’s likely responses to change, both in terms of single-species extinction and system-wide collapse.

## 1 Model and Methods

### 1.1 Mapping system responses from strict biodiversity stability to collapse

To predict an ecosystem’s resilience as well as structural stability we locate boundaries and regimes in parameter-space (including standard deviation of interaction strength, biodiversity and connectivity) and derive a metric in terms of measurable quantities to place the system within this space. To capture responses we analyse large GeneralizedLotka-Volterra (GLV) models

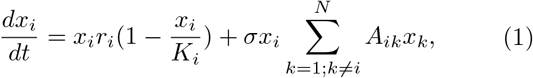

where *N* is the number of species, *x*_*i*_ are species abundances and, *r*_*i*_ and *K*_*i*_ are the intrinsic growth rates and carrying capacities for each species *i. A* is a weighted *N* × *N* adjacency matrix that encodes the interaction strengths (fluxes of bio-masses and other materials or processes) between all species (except intra-specific interactions) and *A*_*ij*_ the specific strength of how species *i* is affected by species *j*. The connectance, *c*, is the probability that an off-diagonal entry of *A* is non-zero, and we sample interaction strengths from a normal distribution with mean *μ* = 0 and a variance of 1. The absence of structure in the interaction matrix is chosen as a starting point when extending the analysis to include the different aspects of stability. We believe that the conclusions carry over to more biologically structured systems. Preliminary investigations confirm this, but a more rigorous investigation is saved for future work. All diagonal entries of *A* are zero and *σ* is a parameter that tunes the standard deviation of inter-specific interaction strengths. Setting *K*_*i*_ = *r*_*i*_ = 1 for all species retrieves the interaction matrix corresponding to the Jacobian used by May (*σ A* − *I*) also inheriting the fact that when *σ* = 0 this represents a system with only self-competitive and isolated primary producers. The fixed-point abundances 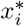 of the system are

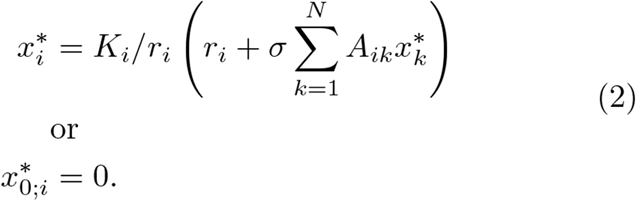

Note that the fixed-points can include species with zero abundance, interpreted as species extinctions. A fixed point without extinct species is called feasible. By allowing for extinctions we explicitly include persistence (fraction of non-extinct species) when mapping the region of resilience (defined as local stability of the fixed-point). This means that *N* is the initial biodiversity whereas the actual biodiversity is the number of species, *n*, with positive abundance at the fixed-point at the time of measurement. In Fig. 1a we show how the fixed-points for a specific system with fixed interactions and parameters (*A*, *c*, *μ*, *K*, and *r*) change as *σ* increases. Importantly, single-species extinctions occur as a response to increased complexity (measured here by *σ*) well before the complexity boundary introduced by May.

**Figure 1.**
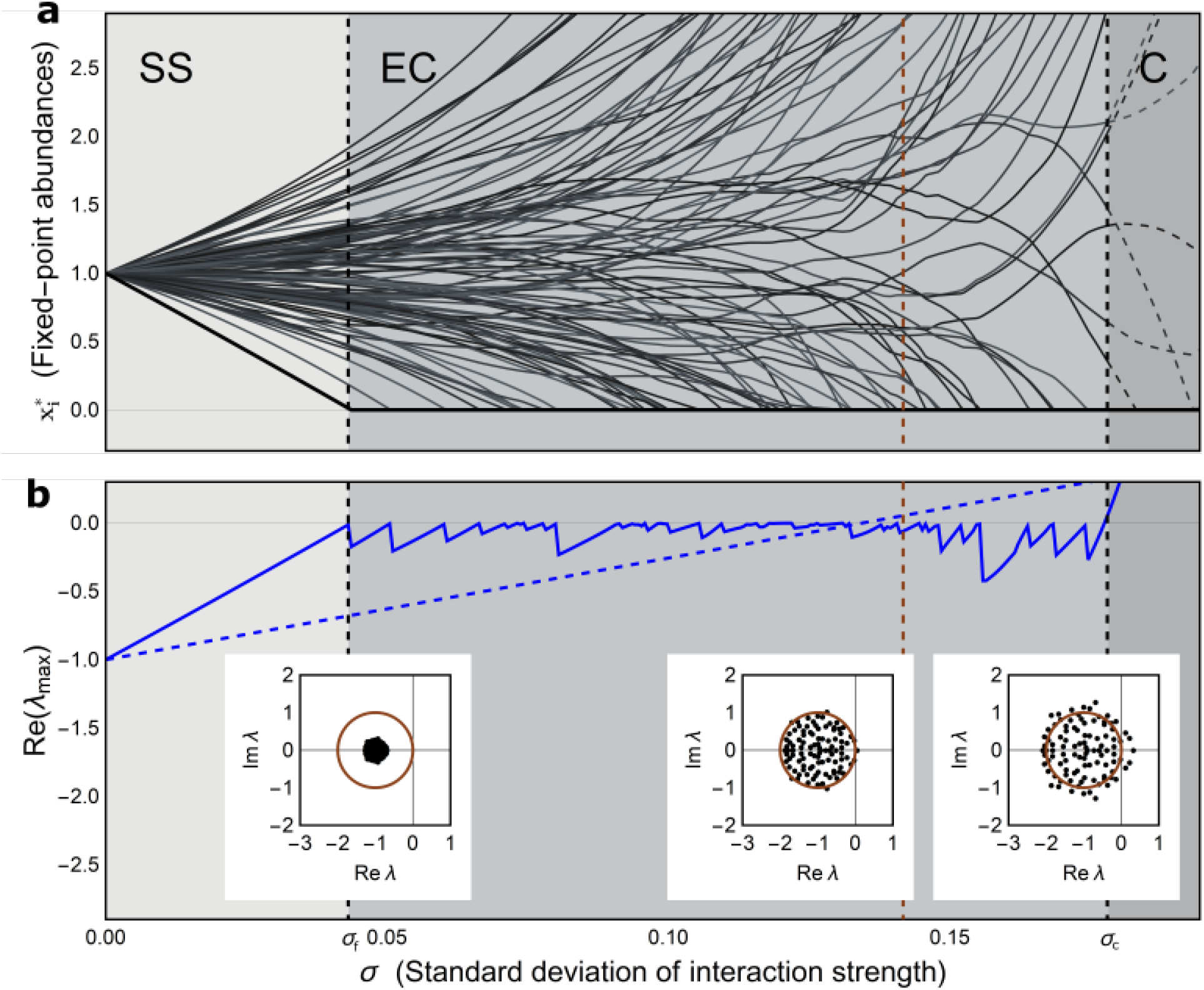
Effects on stability of increasing interaction strength in a complex system. (**a,b**) Example simulation of a system with initial biodiversity, *N* = 100, connectance (fraction of realised species interactions), *c* = 0.5, *r_i_* = *K_i_* = 1, and *μ* = 0 for the mean of the distribution of inter-specific interaction. The plot shows the species abundances (**a**) and the eigenvalue with smallest negative real part (**b**) at resilient fixed-points for increasing values of the standard deviation of interaction strength, *σ*. Three phases of system behaviour, Strict Stability (SS), Extinction Continuum (EC), and Collapse (C) are indicated by the different shades of grey. The first extinction and collapse boundaries *σ*_*f*_ and *σ*_*c*_ respectively are indicated by the dashed black lines. Up to the first extinction, the system is in a feasible (all *N* = 100 species are non-extinct, 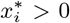), resilient, and structurally stable fixed-point. After the first extinction, the system enters a phase of dynamic self-regulation of complexity via successive single-species extinction, where it is resilient but structurally unstable. This phase includes the complexity limit introduced by May for the initial biodiversity (brown dashed line) and ends where the fixed-points lose resilience altogether. The eigenvalue with smallest negative real part clearly displays the difference in stability behaviour between analysis including and excluding extinctions (**b**). The eigenvalue with smallest negative real part of the reduced community matrix (blue line) fluctuates below zero due to single-species extinctions, while the eigenvalue with smallest negative real part in May’s framework without extinctions (blue dashed line) increases linearly and passes through zero at 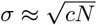. For further clarification, the insets show the eigenvalues for May’s model at the transition points of the system, the brown circle indicating the maximum radius of resilience.

When analysing these systems we are tracking the same fixed-point while increasing *σ*. The resilience is determined by the eigenvalues of the community matrix at this fixedpoint (mathematically equivalent to the Jacobian matrix *J*\*). We keep only non-extinct species in the stability analysis and the resulting reduced community matrix is

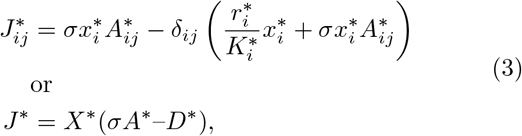

where the superscript asterisk means that we only include non-extinct species. *A*\* is a reduced interaction matrix. The *X*\* and *D*\* are diagonal *n* × *n* matrices with 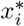 and 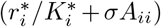 on the diagonal respectively. *A*_*ii*_ is zero for all *i*, and could be omitted but is included for clarity. The *δ* represents the identity matrix, i.e. *δ*_*ij*_ equal to one when *i* = *j* and zero otherwise. We treat the zero abundances at the fixed-point as extinctions. In SI Sec. 1, we discuss systems in which species have a chance to re-invade [41, 42].

For resilience the real parts of all eigenvalues must be negative. When at least one eigenvalue has a positive real part the system is unstable because a change in species abundance caused by a small disturbance will increase exponentially. Changes in parameters (e.g. *σ*) that cause the real part of the eigenvalue with the largest real part to approach zero are usually considered to cause destabilisation, often interpreted as a preamble to collapse. Mathematically, however, collapse is not necessarily implied.

Indeed, in the Extinction Continuum the eigenvalue with smallest negative real part fluctuates at or just below zero (Fig. 1b). The key observation is that single-species extinctions help the system to stay resilient and prevent collapse. Extinctions and stability aspects of the GLV model are demonstrated in Fig. 1. The system response divides into three distinct phases while increasing *σ*. The first phase, Strict Stability (SS), is characterised by a fixed-point that is resilient, structurally stable and feasible (including the full initial biodiversity). The largest real part out of the eigenvalues is negative but approaches zero from below as *σ* increases, at the same rate and magnitude as the smallest species abundance approaches zero [43]. The boundary of the SS phase is located where the eigenvalue with smallest negative real part reaches zero and the first single-species extinction occurs.

In the second phase, Extinction Continuum (EC), the system can uphold resilience with the eigenvalue with smallest negative real part below zero, by single-species extinctions. However, because of extinctions the system is structurally unstable and with only a subset *n* of the initial species *N*. These extinctions occur when *σ* increases, but also for decreasing *σ*, discussed further in SI Sec. 2. This implies that perturbations of systems in this region typically cause extinctions but not a system-wide disruption.

In the third phase, Collapse (C), no stable fixed-point without radical system change is found, and the system’s behaviour is unpredictable. There is the possibility of limit cycles, chaotic behaviour, or a substantial decrease in the number of viable species, resulting in a biodiversity well below the level at the collapse-boundary. This implies that systems are structurally unstable until the collapse-boundary, but they typically undergo more radical qualitative changes than single-species extinctions if pushed to higher complexities.

Note that these three phases are present for the majority of systems with generically chosen intrinsic growth rates *r*_*i*_. The EC phase can always be eliminated if growth rates and/or carrying capacities are specifically chosen for such a purpose for a specific *A*, but such choices violate the assumption of randomness.

With Fig. 1 in mind it is clear that assuming feasibility [16, 26, 31, 36, 44] amounts to incorrect predictions of collapse since it ignores single-species extinctions as a stabilising mechanism (upholding resilience). Or stated differently, for random systems the parameter region of feasibility is a subset of the region of resilience as noted by [43] for competitive/mutualistic systems, [39] including cascade and niche structured food-webs and [45] showing the universality of the feasibility region. The boundary May introduced states that a random system is stable when 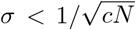 which implies that fixed-points are unstable for 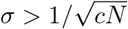. This boundary based on the initial biodiversity actually falls within the EC phase as seen in Fig. 1 and hence does not predict collapse.

It is interesting to note however, that the actual collapse occurs when the reduced system is at 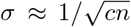 (with a slight upward bias), corresponding to an effective version of the complexity boundary. This further implies that the prediction still holds even though the systems that collapse at this boundary no longer have uncorrelated species interactions (SI Sec. 3 argues for the non-randomness of systems in the EC). Instead, the systems have been dynamically pruned in accordance to dynamics dictated by growth rates *r*_*i*_ and carrying capacities *K*_*i*_. Out of all possible and reasonable values and combinations of *r*_*i*_ and *K*_*i*_, only a small number lead to resilient communities that stay structurally stable for large *σ*. Moreover, that number decreases exponentially as system size increases [45].

### 1.2 Boundaries and persistence

To construct a new metric for predicting collapse, we derive expressions for the two boundaries surrounding the Extinction Continuum as well as persistence. The first boundary between the SS and EC phases is marked by the first extinction event, also corresponding to when the largest real part out of the eigenvalues hits but does not pass zero (since the system size is reduced). To locate this boundary we analyse the interaction matrix *A* using order statistics. This gives a direct estimate of the first-extinction event rather than implicit methods [46].

By writing the fixed-point solutions of Eq. 2 in linear form it can be seen that the species abundances are weighted sums of the entries of *A* (for full derivation see SI Sec. 2). As sums of random variables, the species abundances can themselves be interpreted as random variables. They are found to be distributed according to a normal distribution truncated at zero

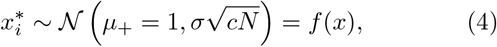

where *μ*_+_ = 1 depends on the choice *r*_*i*_ = *K*_*i*_ = 1 discussed further in SI Sec. 4.2. With this set of species abundance random variables, we can use order statistics to obtain an estimate of the smallest abundance in the set, in a similar manner as [22, 43]. From order statistics the distribution of the minimum (*f_min_*(*x*)) of a set of *N* random variables (in our case 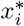) distributed according to *f* (*x*) is

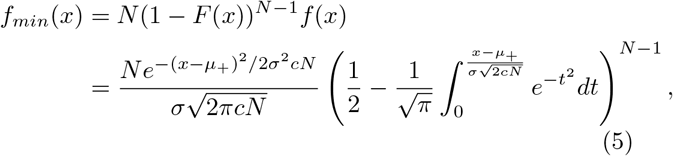

where *F* (*x*) is the cumulative distribution function of *f* (*x*). The first extinction event is at the *σ*_*f*_ for which the mean of the above distribution is zero.

To locate the second boundary between Extinction Continuum and Collapse we make use of a prediction of the persistence in combination with the complexity boundary introduced by May. We predict the persistence from a reduced interaction matrix, shown in Fig. 2 (full derivation SI Sec. 5). We also know that the complexity boundary introduced by May is a good predictor of collapse if all species are guaranteed to be non-extinct [44]. With these two estimates we can locate the collapse boundary for the full system

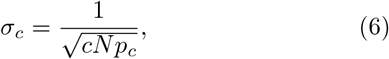

where *p_c_* is the persistence at the collapse boundary (similar *p_c_* found in [40]).

**Figure 2.**
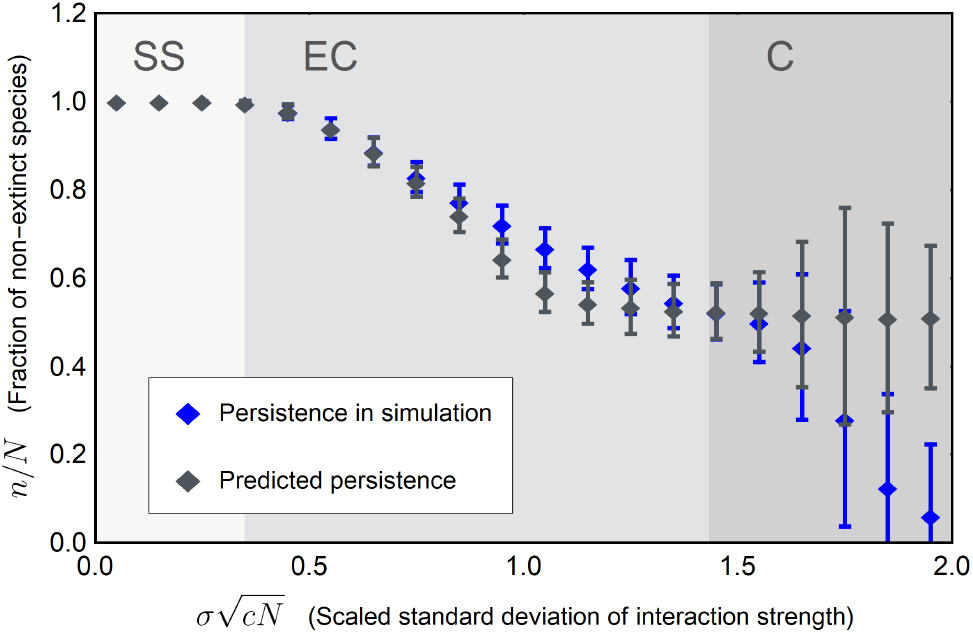
Predicted persistence. The plot shows the average persistence from simulations of systems ranging from *N* = 20 to *N* = 1000 (blue dots) with one standard deviation errorbars and predicted persistence for randomly generated interaction matrices *A* in the same system size range (grey dots) with one standard deviation errorbars. The background shading indicates the three phases of behaviour with the boundaries at the theoretical predictions. The difference between predicted persistence and simulated persistence in the Collapse (C) phase is because when simulating we set a system’s size to zero after collapse. Including these causes the statistics of the fraction *n/N* for the simulated systems to tend to zero. Before this artificial drop towards zero in the simulated fraction in the C phase we can predict the persistence well in the Extinction Continuum.

The results above show that, when the initial biodiversity is known, we can predict the two boundaries and approximate rate of extinctions in the Extinction Continuum for systems of all sizes, as seen in Fig. 3.

**Figure 3.**
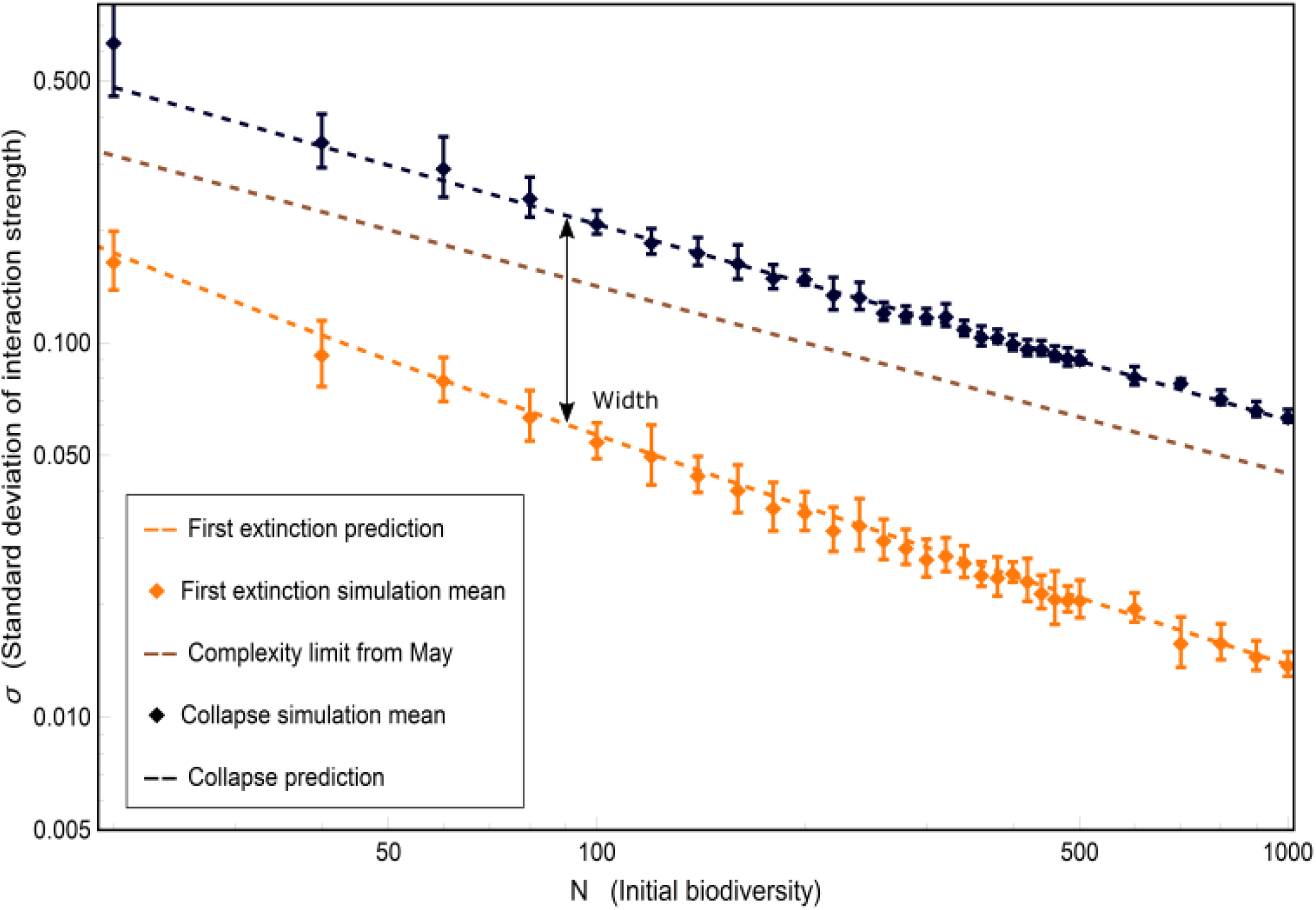
Stability predictions for complex systems. In the parameter-space of the standard deviation of interaction strength, there are three phases of behaviour: Strict Stability (SS), Extinction Continuum (EC), and Collapse (C). Here we demonstrate that these phases hold across a large range of system sizes *N*. The plot shows simulation averages of first extinction events (orange dots) with one standard deviation error bars, our theoretical prediction of first extinction (orange dashed line), the complexity limit introduced by May (brown line), simulation averages of collapse (black dots) with one standard deviation error bars, and our theoretical collapse prediction (black dashed line). The width of the Extinction Continuum is indicated by the arrow, note the increase in width for larger systems. All simulations shown were run with, *r_i_* = *K_i_* = 1, *μ* = 0 for the distribution of inter-specific interactions and a value of *c* = 0.5 for connectance in the interaction matrix *A*.

### 1.3 Instability metric

As stated, if we know the initial biodiversity *N*, we can predict the distance in *σ*-space to collapse for a system of any size whether in the SS or EC phase, since the first extinction boundary *σ*_*f*_ and collapse boundary *σ_c_* are functions of *N*. A system with standard deviation of interaction strength greater than needed for the first extinction boundary (*σ* > *σ*_*f*_) is situated in the Extinction Continuum. This means the biodiversity *n* cannot equal the initial biodiversity *N*. Since *N* cannot be measured for real systems we estimate it from the first extinction prediction *σ*(*n*) and the persistence approximation. With an estimated initial biodiversity *N*_*pred*_, we can properly place the system in parameter-space and calculate its proximity to collapse.

A first thing to note is that a system with *σ* < *σ*_*f*_ (*n*) can be located in the SS phase (if *σ* is smaller but *σ* ≈ *σ*_*f*_ the system might be in EC). If on the other hand *σ* ≈ *σ*_*f*_ (*n*) or *σ* > *σ*_*f*_ (*n*) the system likely resides in the EC phase, and we need to approximate the size of the initial biodiversity. Again, we use the reduced interaction matrix *A*\* to obtain an average rate of extinctions for increasing *σ*. We use this rate to extrapolate backwards to *σ*_*f*_ for the approximation *N*_*pred*_ (more details in SI Sec. 6).

With the approximate initial biodiversity, we predict the two boundaries for the system and construct the metric *γ* to quantify the proximity to collapse

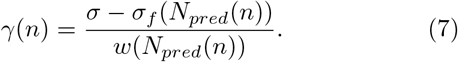

Where *w*(*N*_*pred*_(*n*)) is the width of the EC, *σ*_*c*_(*N*_*pred*_(*n*)) − *σ*_*f*_ (*N*_*pred*_(*n*)). The metric *γ* is defined only in the EC and is a real number [0, 1], although the boundaries of this interval are not exact since we are inferring from the biodiversity, *n*. This metric quantifies the structural instability of the system. We posit that the higher the value of *γ*, the smaller the perturbation that will lead to single-species extinctions and the more probable that perturbations or external pressure will lead to collapse.

## 2 Results

### 2.1 Boundary predictions

Our theoretical estimates of first extinction and boundary to collapse along with the complexity limit introduced by May compared to simulation averages are shown in Fig. 3. The analytic estimates are in good agreement with the results of numerical simulations.

The predictions hold for systems with random interactions between species, interaction strengths sampled from a normal distribution with a mean of zero and variance of one, and different sizes as shown in Fig. 3. In addition, in Figs. 6-10 in SI Sec. 7 we show that there are no qualitative differences in results even if any of the assumptions about distribution, structure, mean, *r*_*i*_, and *K*_*i*_ respectively are modified. The robustness of the existence of the Extinction Continuum follows from the fact that there are exponentially many fixed points that the system can switch between to uphold resilience. Indeed, for this reason we expect the Extinction Continuum to exist in more general population models than the GLV.

### 2.2 Proximity to collapse

To evaluate the *γ* metric we simulated systems with initial biodiversity that ranged from *N* = 20 to *N* = 1000 for values of *σ* placing them in the Extinction Continuum. We calculated *γ* using information about the non-extinct species: the interaction matrix, intrinsic growth rates, carrying capacities, and biodiversity. The predicted *γ* from these simulations are shown in Fig. 4 in comparison to the actual *γ_sim_* from simulations.

**Figure 4.**
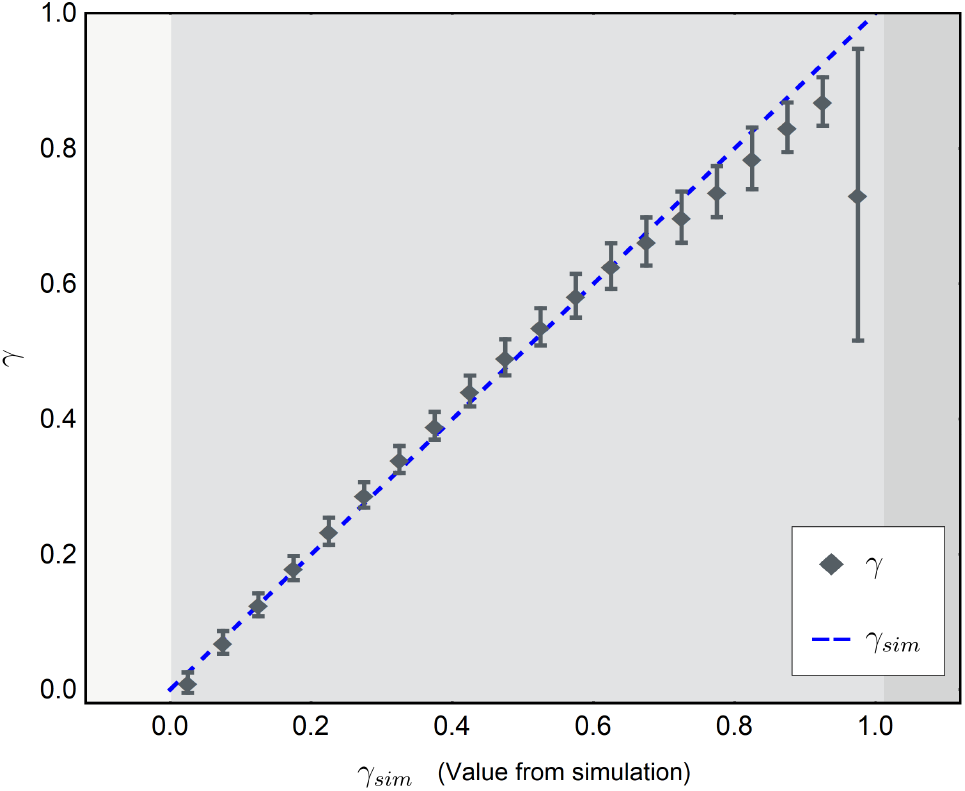
Metric of stability of a complex system. The plot shows the *γ* (dark dots) with one standard deviation errorbars compared to *γ*_*sim*_ (blue dashed line). The *γ*’s were calculated from simulations ranging between *N* = 20 and *N* = 1000 averaged over 10 replicates. The background shading indicates the three phases of behaviour. Here we see that our metric of proximity to collapse is within one standard deviation of the correct value right up until collapse, which means it is a good predictor of position in parameter-space.

As seen in Fig. 4, *γ* follows the actual *γ_sim_* closely. The variance of the prediction increases during the EC as expected, but the actual value is within one standard deviation right up until collapse.

All simulations for Fig. 4 were performed for a standard setup with *r*_*i*_ = *K*_*i*_ = 1, no symmetries in the interaction matrix *A*, and the interactions strengths drawn from a Normal distribution with zero mean. For small deviations of the mean from zero (|*μ*| < 0.05), the *γ* metric remains a reliable approximation. For larger deviations the prediction of collapse becomes more complicated. This is due to a larger impact on stability of the species abundances 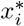 and that we use the general (*σA*\* − *I*) instead of the actual Jacobian of the system *X*\*(*σA*\* − *I*) when predicting persistence (since the 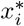 are unknown). This problem is shared with the complexity boundary introduced by May and discussed further in SI Sec. 8.

As an additional verification that *γ* quantifies proximity to collapse, we gathered statistics for resilience of reduced systems from the entire Extinction Continuum before and after perturbations of its interaction matrix. The perturbation was an addition of Gaussian noise (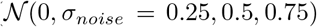) to all the realised interactions in the interaction matrix. This perturbs the variance in the interaction strength distribution and relative interaction strengths but leaves the connectance and topology of interactions unaltered as well as the intraspecific interactions.

The results from these simulations can be seen in Fig. 5. The top panel shows the percentage of systems that find a resilient fixed-point after the perturbation and the bottom panel shows the relative size of the resilient communities after the perturbation. The first thing to note is that the resilient communities after perturbation are consistently of smaller size (but have unchanged connectance SI Sec. 3), even for small perturbations Fig. 5b. It is also evident that systems with larger *γ* are closer to collapse and lose more species before finding a resilient state, a sign of increasing structural instability. This is consistent with our previous observation that correlations are introduced in the dynamic reduction of systems needed for resilience. It is also clear from Fig. 5a that systems with a large *γ* are more likely to collapse after a perturbation. Both these findings show *γ* is a metric for collapse.

**Figure 5.**
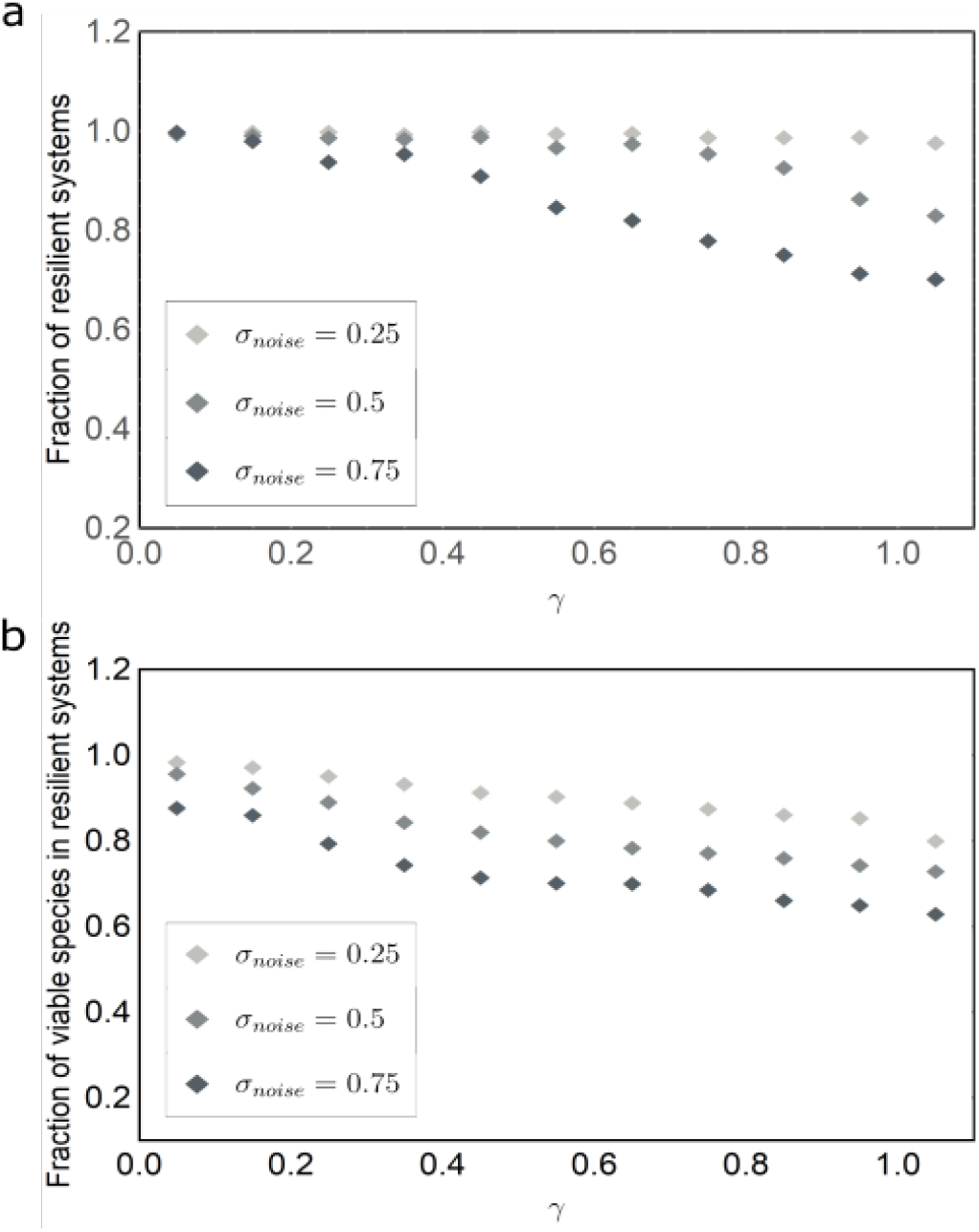
Predicting collapse. The plot shows in the top panel the fraction of systems for a certain *γ* that found a resilient fixed point after perturbations. The systems with varying *γ* values were generated from random systems with *N* = 70 and *N* = 100, connectance *c* = 0.5, intrinsic growth rates and carrying capacities *r_i_* = *K_i_* = 1, and interaction strengths from a Normal distribution with *μ* = 0, by specifying their standard deviation of interaction strength *σ*. Note that *γ* reaches values larger than one, this is because it is inferred from *N*_*pred*_ simulating a situation where the initial biodiversity is unknown. The bottom panel shows the fraction of non-extinct species at the new fixed-point for systems that found a resilient state after perturbation. Here even for small perturbations in the Extinction Continuum some species go extinct for the system to find a new resilient fixed-point. This effect increases for larger *γ* indicating more specificity of interactions in more dynamically-pruned systems. Together the plots demonstrate that a larger *γ* indicates collapse both in terms of a substantial loss of species (more structurally unstable) and a higher probability of loss of resilience.

## 3 Discussion and conclusions

We have investigated the conditions for which an ecosystem is resilient and structurally stable. In addition, we derived a metric that uses only measurable quantities and that quantifies the proximity to collapse and level of structural stability. It has been noted in various studies that both feasibility and local stability are important concepts for ecosystems [39, 40, 43, 47] and that real systems tend to grow in size to the point at which they are structurally unstable [48]. By combining resilience, structural stability, and persistence, we show that there exists parameter ranges for which a system can be resilient but structurally unstable, an observation that introduces a previously overlooked phase in parameter space. It also shows that for a system with random interactions, feasibility is lost at lower levels of complexity than is resilience, which means systems with complexity levels above the first extinction boundary typically need non-random interaction patterns in order to be resilient.

The Extinction Continuum and the accompanying behaviour of the eigenvalue with smallest negative real part is relevant when trying to predict collapse with critical slowdown – a system starts to recover very slowly after perturbations in species abundances – the standard method for proximity detection [49–51]. In our analysis systems show critical slowdown throughout the EC phase, but the slowdown only involves one or a few species at the time (the species that will go extinct next) as in [52]. The major transition and predictor of proximity to system collapse however is when the system is close to the collapse-boundary and the critical slowdown involves many species at once. In contrast to critical slowdown our metric can be used to better predict the location of the collapse early in the Extinction Continuum, when the system is still far away from the collapsing point and show no sign of system-wide slowdown.

It has been argued that all real systems self-organise to structural instability [48]. The rationale behind this is that there can always be immigration, adaptation or speciation until the point where additional species will cause others to go extinct. This will stabilise the community at a certain biodiversity but with species turn-over. The idea implies that a system self-organises to the largest system theoretically possible [40, 53], which would be at the complexity boundary introduced by May (and its extensions from later studies taking interaction structures into account [36, 44]). This would be the case if there is a large enough regional species-pool from which a variety of species can immigrate or long enough time for multiple speciations. Here, rather than a boundary, we have shown that there is an entirely new phase of structural instability. As we have shown, for communities more complex than the first extinction boundary, correlations are needed, which reduce the number of species that have the ability to invade. In case of a less diverse species-pool, it is possible that systems will stabilise in the Extinction Continuum. This might even be likely since empirical studies have found no correspondence with May’s limit [54]. Also, in line with the hypothesis that systems will evolve to maximise structural stability [37] systems in the EC phase are structurally unstable but less so when further from the collapse boundary.

We argue that all systems beyond the first extinction boundary need finely tuned, or perhaps naturally selected, parameters to be resilient. A system can be feasible at the collapse-boundary for certain growth rates and/or carrying capacities [45]. Since in our analysis growth rates and carrying capacities are fixed (or drawn from a distribution), a system can only stay resilient if correlations are introduced through the dynamics by non-random species extinctions. These correlations can either be within the reduced interaction matrix *A*\* or between *A*\* and the fixed-point species abundances. The latter having the largest stabilising effect [55]. Correlations within *A*\* were found to be positive for 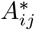 and 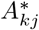 and negative for 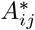 and 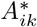 [55]. This suggests that a species having either a positive or negative effect on the community in general and species that can balance negative encounters (for example plants competing for sunlight) with positive encounters (for example seed dispersal) with other species are stabilising for a community. For predator/prey systems we found negative correlations for both 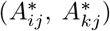 and 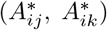 hinting at secondary consumers as stabilising for food-webs. Although this might be too strong a conclusion since there are no trophic hierarchies present. It has also been found that correlations in interaction strengths weighted by biomass (between 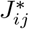 and 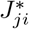) increase stability (in terms of increasing the range of resilience) [13]. Even though this might lead to a larger range of resilience, as we have shown the correlations introduced by the dynamics still make the systems structurally unstable.

The *γ* metric can be calculated from observable quantities of an ecosystem that capture proximity to collapse. Since changes in the GLV parameters (*N*, *c*, *r*_*i*_, *K*_*i*_, distribution of interaction strengths, and symmetries in interactions) induce “mere” shifts in boundaries we argue that *γ* not only quantifies likelihood of collapse in terms of *σ* perturbations but also structural stability in general. This makes *γ* a more broadly applicable metric than its definition (Eq. 7) might indicate. Inherently it also elucidates system response behaviour beyond the feasibility region, in addition to the feasibility investigation in [39], with extinctions acting as a stabilising mechanism. In [45] a different way of capturing structural stability is proposed, measured as the size and shape of the feasibility region of a system when varying intrinsic growth rates. Although their approach requires a slightly stricter resilience criteria excluding some interaction matrices, in line with *γ* their results lead to decreasing structural stability with increasing *σ*. The structural stability measures also differ in that *γ* can be evaluated for a specific system while in [45] the measure applies to systems with unknown intrinsic growth rates.

For the *γ*-metric, in addition to the GLV parameters, estimates are needed for the interaction strengths for each interaction in the matrix. Metabolic theory may help guide this based on the size and temperature of the species involved [56, 57]. Note however that we include all types of biological interactions. This is in contrast to many previous studies looking for stability criteria of ecosystems that independently focus on food-webs [3, 11, 28] or different nontrophic interactions [42, 46, 58]. There is always a risk of feed-backs between these different aspects that may undermine the results. Studies have highlighted such feed-backs [25, 59], and encouraging steps are being taken towards multilayer representations of ecosystems [27]. Thus for *γ* to be a more realistic measure it remains to be shown in future studies if it can be extended to a multi-layer framework instead of having all types of interactions in a single layer in the interaction matrix.

Even though we posit a wide scope for *γ* there are important limitations of our current study. Apart from symmetric (mutualistic/competitive) and anti-symmetric (predator/prey) interactions, we have not investigated the effects of structured interaction topologies, that might expand the feasibility region [39]. Introducing more realistic structures such as trophic hierarchies into the interaction matrix is an important direction for future work. Another limitation to our approach, and the GLV approach in general, is that the interaction matrices are assumed to be rigid – static “averages” of interactions among species [60–62]. In reality interactions might fluctuate with external factors such as seasons or change due to adaptive foraging behaviour. Since seasonal changes in general do not lead to extinctions, it is assumed that they do no not introduce fluctuations large enough to breach the limits of structural stability for systems in the Extinction Continuum. Still, we do not know, for example, if cycles of interaction strengths play a role in the stability of communities. Studies with adaptive foragers have found that they can be either stabilising (in terms of structural stability) [15] or sometimes destabilising (in terms of secondary extinctions) depending on timescale when compared to rigid systems [17]. Since adaptive foraging may affect structural stability positively but does not exclude secondary extinctions, it is unclear how the EC would be affected by changing interactions.

In conclusion we have developed a theoretical analysis to quantify a system’s proximity to collapse based on measurable information such as biodiversity, connectance, and species interactions strengths. The *γ* metric is both detectable at larger parameter-”distances” from collapse and easier to evaluate as compared to, for example, critical slowdown. We also present a consistent framework that includes resilience, structural stability, and persistence, in which to apply the *γ* metric. Together the framework and *γ* metric expand our ability to estimate a system’s likely response to perturbations and assess its risk of collapse.

## Author Contributions

Conceived the project: M.N.J.; Performed simulations and wrote paper: S.P.; Analytic results: M.N.J., S.P.; Interpretation of results and comments on manuscript: M.N.J., S.P., V.M.S.

## Author Information

The authors declare no competing financial interests.

