## Supplementary information for "Predicting Collapse of Complex Ecological Systems: Quantifying the Stability-Complexity Continuum"

November 21, 2019

#### Contents

**1 Initial species biodiversity or species-pool** **2**

**2 Extinctions when decreasing complexity** **2**

**3 Correlations in the reduced interaction matrices in the Extinction Continuum** **4**

**4 First extinction event** **6**

4.1 When $r_i = K_i = 1$ . . . . . . . . . . 6

4.2 The general case of intrinsic growth rates and carrying capacities $r$ and $K$ . . . . . . . . . 7

**5 Collapse boundary and persistence** **8**

**6 Estimating initial biodiversity $N_{pred}$** **9**

**7 The extinction Continuum persists** **9**

**8 The case of large distribution mean $|\mu|$** **15**

#### 1 Initial species biodiversity or species-pool

In our analysis we treat species abundances equal to zero as "complete" extinctions and not only as local phenomena. This is in contrast to other studies which treat the initial biodiversity rather as a species-pool (species present in a region consisting of smaller local communities). In the species-pool interpretation if a species goes extinct in a local community it is not lost and it can still re-invade the local community at a later point in time [1, 2]. Specifically in [1] they found different phases of local community behaviour/turn-over when increasing the mean and variance of interaction strengths. In general increasing turn-over for increasing variance and mean of interaction strength.

The full Jacobian of the system is

$$J_{ij} = \delta_{ij} \left( r_i - 2\frac{r_i}{K_i} x_i + \sigma \sum_{k=1;k\neq i}^{N} A_{ik} x_k - \sigma x_i A_{ij} \right) + \sigma x_i A_{ij}, \tag{1}$$

where $r_i$ and $K_i$ are intrinsic growth rates and carrying capacities respectively, $A$ the interaction matrix with zero diagonal (hence $-\delta_{ij}\sigma x_i A_{ij} = 0$ in Eq. 1 and can be excluded), $x_i$ species abundances, $\sigma$ the standard deviation of interaction strengths, and $\delta$ the Kronecker delta (equal to one when $i = j$ and zero otherwise). A reduced Jacobian $J^*$ for only viable (non-extinct) species, with fixed-point solutions

$$x_i^* = K_i^*/r_i^* \left( r_i^* + \sigma \sum_{k=1;k\neq i}^{n} A_{ik}^* x_k^* \right), \tag{2}$$

is given by

$$\begin{aligned} J_{ij}^* &= \sigma x_i^* A_{ij}^* - \delta_{ij} \left( \frac{r_i^*}{K_i^*} x_i^* + \sigma x_i^* A_{ij}^* \right) \\ &\text{or} \\ J^* &= X^*(\sigma A^* - D^*), \end{aligned} \tag{3}$$

where $X^*$ and $D^*$ in the linear algebra formulation are diagonal matrices with species fixed-point abundances and $(r_i^*/K_i^* + \sigma A_{ii}^*)$ on the diagonal respectively, although $A_{ii}^* = 0$ for all $i$. The superscript asterisks indicate the inclusion of only non-extinct viable species. This is the community matrix (Jacobian at a fixed point) which we use in our stability analysis.

At a fixed-point including extinct species the Jacobian for an extinct species $i$ would be

$$J_{ij}^{x_i^*=0} = \delta_{ij} \left( r_i + \sigma \sum_{k=1;k\neq i}^{N} A_{ik} x_k^* \right). \tag{4}$$

Since this is a diagonal matrix it shows the eigenvalues for the extinct species. For small standard deviation of interaction strengths $\sigma$, the eigenvalues are generally negative, hence stable. For larger $\sigma$ this is not generally the case indicating that the extinct species could re-invade, were they not completely extinct. Thus in the latter part of the Extinction Continuum there can be multiple stable fixed-points in accordance with [2, 3] with different combinations of viable species from a species-pool including "already" extinct ones. Although their basins of attraction differ in size. With no re-invasion on the other hand, larger $\sigma$ hence less structurally stable systems means a risk of substantial biodiversity reduction if system parameters change (such as $r_i$ or $\sigma$), corresponding to the larger turn-over (switching of stable fixed-points) for larger $\sigma$ found in [1]. The difference then in the two interpretations, complete or local extinctions, is that if system parameters change the structural instability lead to either decreased biodiversity (complete) or turn-over while sustaining biodiversity (local).

#### 2 Extinctions when decreasing complexity

One interesting and perhaps counter intuitive result is that extinctions can occur in the Extinction Continuum as a response to decreasing the standard deviation of interaction strength $\sigma$, corresponding to decreasing complexity. We tested this for systems with random interaction matrices but with intrinsic growth rates $r_i$ specifically chosen to make them resilient and feasible at the complexity boundary introduced by May $\frac{1}{\sqrt{cN}}$. This was done both by brute force sifting through vectors $\mathbf{r}$ with non-negative $r_i$ drawn from truncated Normal and Uniform distributions centred at 1, or allowing for negative $r_i$, by $\mathbf{r} = (I - \sigma A)\mathbf{x}_{rand}$, with uniformly distributed $x_{i,rand} > 0$. Indeed, we found systems can exhibit species extinctions when $\sigma$ decreases. This can also occur for our dynamically reduced systems $A^*$ with biodiversity $n$ as seen in Fig. 1.

One interpretation of this response is that for systems to be resilient at levels of complexity high enough to reside in the EC, correlations or interaction patterns have to be in place to accommodate stability. For instance, one species might depend crucially on different species at different levels of complexity, increasing its fragility at these levels if any of the other species go extinct. Thus, correlations seem to be stabilising at specific levels of complexity rather than stabilising for all below the collapse boundary. This makes them sensitive not only to an increase in complexity but also to a decrease, though less so.

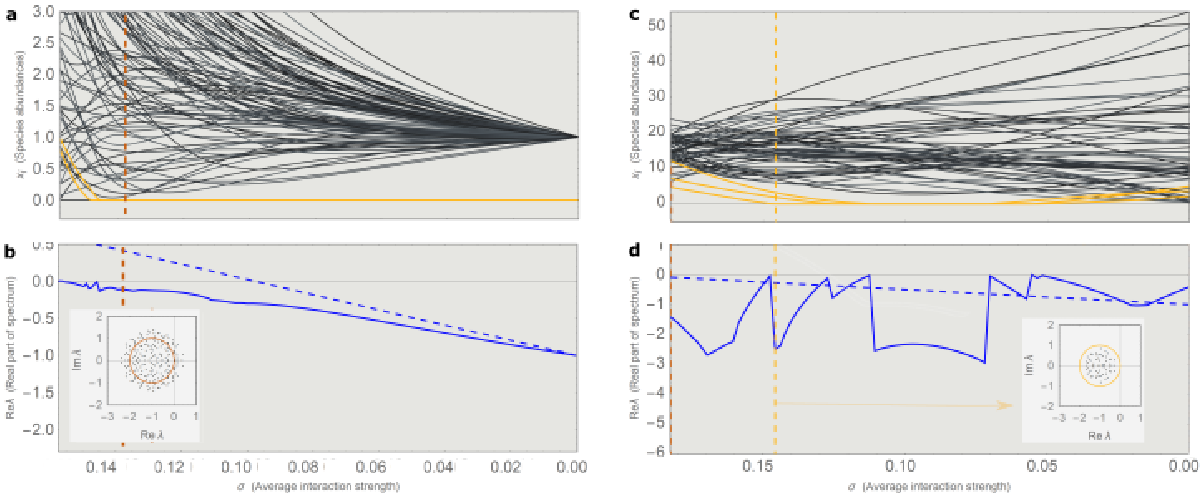


**Figure 1** | **Species extinctions for decreasing $\sigma$.** (**a,b**) An example simulation of a dynamically reduced system with $n = 103$, $c = 0.5$, $K_i = r_i = 1$, and Normally distributed interaction strengths Normal$(0, 1)$, when decreasing $\sigma$. We start the simulation with $\sigma$ equal to the systems collapse value $\sigma_c$ when run with increasing $\sigma$ from zero. The simulation run when increasing $\sigma$ had initial biodiversity $N = 200$ and was reduced to a viable community of $n = 103$ non-extinct species, which is the starting amount in (**a,b**). Figure (**a**) shows that species can go extinct (highlighted in yellow) even when decreasing the interaction strength standard deviation. The leading eigenvalue of the system is shown in (**b**) (blue line) with the leading eigenvalue of May's model (blue dashed line) and the instability prediction of May's model indicated by the brown dashed line. The inset shows the spectrum of May's model at the instability prediction (for $n = 103$) and maximum stability radius (brown circle), note that the extinctions have changed the stability of the system and the spectrum is no longer contained inside the stability radius. (**c,d**) These plots show a system with $N = 60$, $c = 0.5$ and Normal distribution Normal$(0, 1)$ of interaction strengths, where the $r_i$ were chosen such that the system is feasible at the complexity limit introduced by May. Panel (**c**) Shows a run when decreasing $\sigma$ from May's limit. The species going extinct are highlighted in yellow and the first extinction when decreasing $\sigma$ is marked by the yellow dashed line. Panel (**d**) Shows the leading eigenvalue of the system (blue line) and for May's model (blue dashed line). Both are negative the entire range of interaction strengths, but the actual system can only keep stability by extinctions and resurrections (when the leading eigenvalue is seen to hit zero). The inset shows the spectrum of May's model at the first extinction of the actual model (maximum stability radius in yellow), without any sign of instability or impending extinction.

### 3 Correlations in the reduced interaction matrices in the Extinction Continuum

Systems with randomly generated interactions that have parameter values $N$, $c$ and $\sigma$ placing them in the Extinction Continuum must be dynamically pruned through extinctions (or chosen by some other means) in order to be resilient. This pruning (or choice) makes them no longer random. The simplest argument for this is the statistical result of first extinction boundary for random systems. Since random systems with $N$, $c$ ($c$ remains constant in the Extinction Continuum see Fig. 2) cannot keep all $N$ species beyond $\sigma_f$, some non-random feature must come into play. Two observed changes of dynamically pruned systems is the increase of the interaction strength mean as shown in Fig. 3 and the correlations between ($A^*_{ij}$, $A^*_{ik}$) and ($A^*_{ji}$, $A^*_{ki}$) noted by [4] for mixed interactions shown in Fig. 4 together with correlations in predator/prey systems.

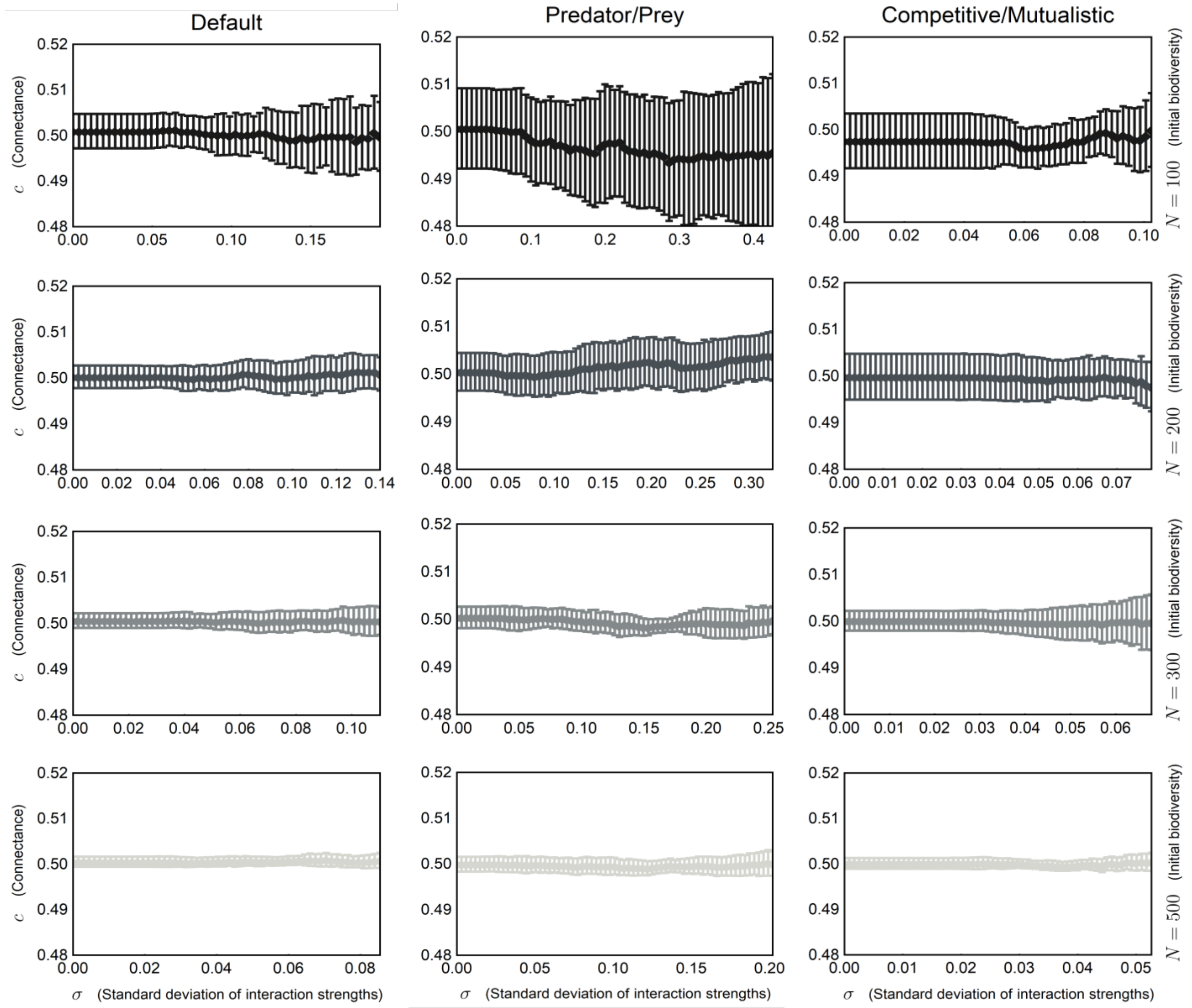


**Figure 2 | Connectance of reduced systems.** The plot shows the mean connectance and one standard deviation errorbars for the connectance the interaction network of dynamically reduced systems as the standard deviation of the interaction strengths $\sigma$ is increased. This is shown for systems with *Default*, *Predator/Prey*, and *Competitive/Mutualistic* structures for initial biodiversities $N = 100, 200, 300, 500$. All systems have intrinsic growth rates and carrying capacities $r_i = K_i = 1$. It is clear that the connectance does not change significantly in the Extinction Continuum.

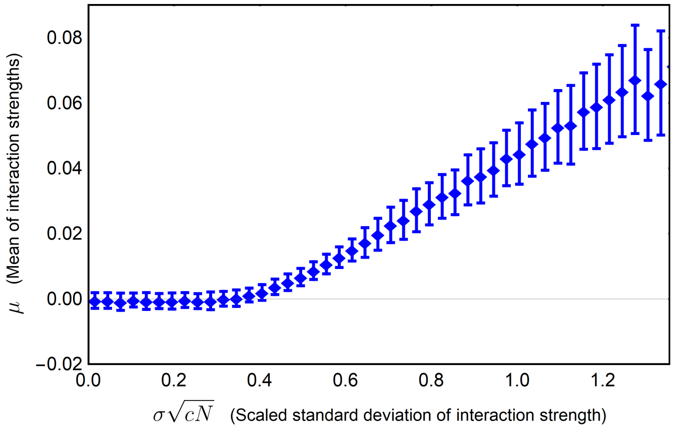


**Figure 3 | Increasing mean when increasing standard deviation of interaction strengths.** The plot shows means and one standard deviation errorbars for the mean of the interaction strengths $\mu$ of dynamically reduced systems as the standard deviation of the interaction strengths $\sigma$ is increased. The systems initial distribution of interaction strengths $A_{ij} \sim \mathcal{N}(0,1)$, intrinsic growth rates and carrying capacities $r_i = K_i = 1$, connectance $c = 0.5$ and initial biodiversity $N$ ranging between 100 and 500 in steps of 20. It is clear that $\mu$ increases in the Extinction Continuum, this is to be expected since species with predominantly negative interactions are expected to go extinct first.

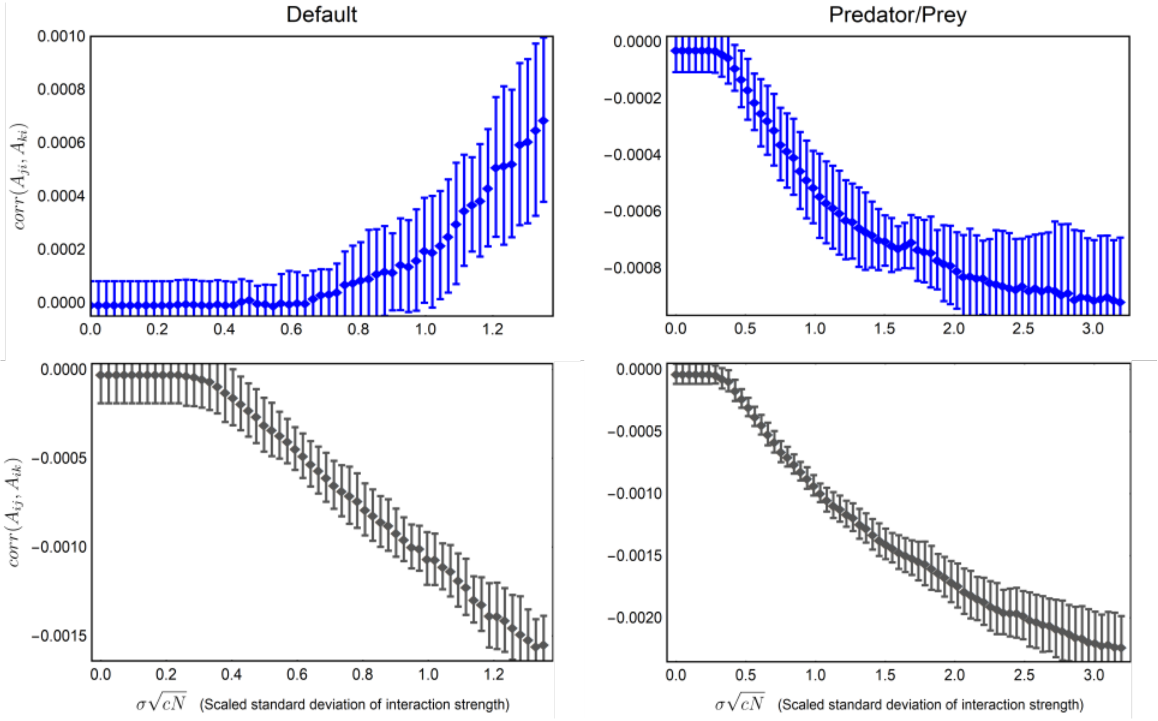


**Figure 4 | Increasing correlations when increasing standard deviation of interaction strengths.** The figure shows correlation means (10 replicates) with one standard deviation errorbars for systems with $A_{ij} \sim \mathcal{N}(0,1)$ in the *Default* column and for systems with $A_{ij} \sim \mathcal{N}(0,1)$ but in addition $\text{sign}(A_{ij}) = -\text{sign}(A_{ji})$ in the *Predator/Prey* column. All systems have connectance $c = 0.5$, intrinsic growth rates and carrying capacities $r_i = K_i = 1$ and initial biodiversity $N = 500$. The correlation patterns are similar for systems of all sizes although the magnitude of correlations decreases for larger systems. Note the switch from positive to negative correlations in $(A^*_{ji}, A^*_{ki})$ in predator/prey systems. This might imply that secondary consumers are stabilising, although this is maybe too strong a conclusion because of the lack of trophic hierarchy in these systems.

### 4 First extinction event

#### 4.1 When $r_i = K_i = 1$

To predict the first extinction boundary we want to find a value of the standard deviation of interaction strength $\sigma_f$ where it is most likely for a system's first extinction event to occur. Because we are interested in the boundary $\sigma_f$ of the Strict Stability (SS) phase we can assume feasible fixed points ($x_i^* > 0$ for all $i = 1, 2 \ldots N$ where $N$ is the initial biodiversity). The fixed points written as linear equations for the case $r_i = K_i = 1$ are

$$(I - \sigma A)\,\mathbf{x}^* = \mathbf{1} \tag{5}$$

where the $\mathbf{1}$ is a vector of ones, $I$ is the identity matrix, $\mathbf{x}^*$ is a vector with fixed point species abundances, and $A$ an $N \times N$ matrix with random interaction strengths from a distribution with mean $\mu$ and variance set to one. The more general case for arbitrary $r_i$ and $K_i$ is given below, supplementary information 4.2. To account for competitive and mutualistic systems with mean interaction strength $\mu \neq 0$ we will include these cases in our analysis. To simplify when $\mu \neq 0$ we separate the interaction matrix into $\mathbf{A} = \mathbf{A}_0 + \mu\mathbf{M}$ where the distribution of the entries in $\mathbf{A}_0$ have mean zero. The diagonal of $A$ is set to zero, the connectance is $c$ and the entries $M_{ij} = 1$ for $A_{ij} \neq 0$. In the case $c = 1$ when all possible interactions are realised, $\mathbf{M}$ is a matrix of ones with zero diagonal. The fixed-point equation can then be written as

$$(\mathbf{I} - \sigma\mathbf{A}_0 - \sigma\mu\mathbf{M})\,\mathbf{x}^* = \mathbf{1}. \tag{6}$$

We begin with the case $c = 1$, so $\mathbf{M}\mathbf{x}^*$ can be approximated with the constant vector $x_{TOT}\mathbf{1}$, with $x_{TOT} = \sum_{i=1}^{N} x_i^*$. The diagonal element is excluded in each sum, but since the variance of the species abundances $x_i^*$ before first extinction is small the approximation is sufficient. The constant vector $x_{TOT}\mathbf{1}$ can be moved to the other side of Eq. 6 and grouped with $\mathbf{1}$. We can then solve for the fixed point abundances

$$\mathbf{x}^* = (\sigma\mu x_{TOT} + 1)\,(I - \sigma\mathbf{A}_0)^{-1}\,\mathbf{1}. \tag{7}$$

For the inverse a von Neumann series expansion ($(\mathbf{I} - \mathbf{B})^{-1} = \sum_{m=0}^{\infty} \mathbf{B}^m$ for some matrix $\mathbf{B}$ with approximately $|\mathbf{B}| < 1$) to first order gives

$$\mathbf{x}^* \approx (\sigma\mu x_{TOT} + 1)\,(I + \sigma\mathbf{A}_0)\,\mathbf{1}. \tag{8}$$

Because the entries of $\mathbf{A}_0$ are drawn from a random distribution, the species abundances $x_i^*$ can be treated as stochastic variables. We thus get a set of $N$ stochastic variables

$$X_i^* \approx (\sigma\mu x_{TOT} + 1)\left(1 + \sigma\sum_{j=1}^{N}(A_0)_{ij}\right). \tag{9}$$

Now that we have estimates of species abundances, we can estimate the smallest $\sigma$ at which there exists a $x_i^* \leq 0$.

Because we assumed $c = 1$, a nonzero $\mu$ only adds a scaling factor to $X_i^*$, hence it will not affect our prediction for when the first species abundance $x_i^*$ becomes zero. Cases when $c < 1$ on the other hand introduce a slight bias. Although we in this case can no longer treat the sum of abundances $\sum_j x_j^*$ in Eq. 8 as a constant $x_{TOT}$ – it is a stochastic variable – it is scaled with the product $\sigma\mu$. When $\sigma\mu$ is small the bias is therefore negligible. Thus, for small $|\mu|$ the scaling factor $x_{TOT}$ for all values of $c$ does not impact our estimate of first extinction.

The distributions of the $X_i^*$ are row sums of the interaction matrix $\mathbf{A}_0$. For a normal distribution of the interaction strengths or any distribution so long as the product of initial biodiversity and connectance is large ($cN >> 1$) enough to invoke the central limit theorem, the sums are normally distributed

$$\begin{aligned} X_i^* &= 1 + \sigma\left((A_0)_{i1} + (A_0)_{i2} + ...(A_0)_{iN}\right) \\ &\sim \mathcal{N}(\mu_+ = 1, \sigma\sqrt{cN}\,), \end{aligned} \tag{10}$$

where $\mu_+$ and $\sigma^2 cN$ are the new means and variances for the sums. The extra $cN$, appearing in the variance is the mean of the Binomial distribution of $N$ trials with success probability equal to the connectance $c$.

From order statistics we get the distribution for the minimum species abundance ($f_{min}(x)$) in the set of abundances from the distribution function of $X_i^*$ ($f(x)$) and its cumulative distribution function, $F(x)$,

$$\begin{aligned} f_{min}(x) &= N(1-F(x))^{N-1}f(x) \\ &= \frac{Ne^{-(x-\mu_+)^2/2\sigma^2 cN}}{\sigma\sqrt{2\pi cN}}\left(\frac{1}{2}-\frac{1}{\sqrt{\pi}}\int_0^{\frac{x-\mu_+}{\sigma\sqrt{2cN}}} e^{-t^2}dt\right)^{N-1} . \end{aligned} \quad (11)$$

The first extinction event is then predicted at the $\sigma_f$ for which the mean of the above distribution is zero. This prediction is in well agreement with simulations as shown in Fig. 2 in the main text.

#### 4.2 The general case of intrinsic growth rates and carrying capacities $r$ and $K$

We have established that the first extinction event for the case when $r_i = K_i = 1$ occurs at the $\sigma$ for which we get a mean of zero for the minimum distribution $f_{min}(x)$ of the set of all species abundances

$$f_{min}(x) = N(1-F(x))^{N-1}f(x). \quad (12)$$

Where the initial biodiversity is $N$, $f(x)$ and $F(x)$ are the distribution function and the cumulative distribution function of the species fixed point abundances $x_i^*$ ($i = 1, 2...N$) respectively. For the more general case of $r_i$ and $K_i$ drawn from distributions the row sums of $A$ used to get the species abundances are now weighted sums by $r_i$ and $K_i$. Thus we need to do more work to capture this extra variability. We begin with the case of $\mu = 0$. The fixed point abundances for the general case written as a linear equation are

$$\begin{aligned} \boldsymbol{x}^* &= \left(\frac{1}{K_i}I - \sigma A\right)^{-1}\mathbf{r} \\ &= K\left(I - \sigma AK\right)^{-1}\mathbf{r}, \end{aligned} \quad (13)$$

where $K$ is a diagonal matrix with $K_i$ on the diagonal. When locating the first extinction event we interpret the fixed point abundances, intrinsic growth rates and carrying capacities as stochastic variables $X_i^*$, $R_i$ and $K_i$. Using a von Neumann expansion to first order for the inverse in equation 13 as outlined above we get an equation for the abundances as

$$X_i^* = K_i\left(R_i + \sum_{j=1}^{N} A_{ij}K_jR_j\right). \quad (14)$$

Here we are assuming a small variance of $K_i$ for the von Neumann expansion to hold. When $r_i$ and $K_i$ are no longer constant but drawn from distributions, $X_i^*$ cannot be approximated by the normal distribution as before. This is because we add and multiply by stochastic variables $R_i$ and $K_i$. A product distribution of two stochastic variables $Z$ and $Y$ with distribution functions $f_Z(z)$ and $f_Y(y)$ respectively, has the form

$$f_W(w) = \int_{-\infty}^{\infty} f_Z(z)f_Y(w/z)\frac{1}{|z|}dz. \quad (15)$$

A sum of two random variables $Z$ and $Y$ has the distribution function

$$f_W(w) = \int_{-\infty}^{\infty} f_Y(w-z)f_Z(z)dz. \quad (16)$$

Combining the three distribution $f_R(r)$, $f_K(k)$, and $f_N(y)$, where the first two are for $R_i$ and $K_i$ respectively, and the last is from the sum $AK\mathbf{r}$ (Eq. 14, where K is a diagonal matrix with $K_i$ on the diagonal), we get

$$f_X(x) = \int_{-\infty}^{\infty}\left(\int_{-\infty}^{\infty} f_N(z-r)f_R(r)dr\right)f_K(x/z)\frac{1}{|z|}dz, \quad (17)$$

with CDF

$$F_X(x) = \int_{-\infty}^{x} \left( \int_{-\infty}^{\infty} \left( \int_{-\infty}^{\infty} f_N(z-r) f_R(r) dr \right) f_K(x/z) \frac{1}{|z|} dz \right) dx. \tag{18}$$

These are the more general functional forms to plug into the minimum distribution in Eq. 12. The mean and variance of the normally distributed sum must also be updated since the row entries of $A$ are now weighted by the product of $R_i$ and $K_i$. The mean and variance of the product of three stochastic variables are

$$\begin{aligned} \mu_{AKR} &= \mu \times \mu_K \times \mu_R = 0 \\ \sigma_{AKR}^2 &= \sigma^2(\sigma_K^2 + \mu_K^2)(\sigma_R^2 + \mu_R^2) - \mu_{AKR}^2, \end{aligned} \tag{19}$$

and since we have the density $c$ and the size of the system $N$, the entire sum is

$$\begin{aligned} \mu_N &= cN\mu_{AKR} = 0 \\ \sigma_N^2 &= cN\ \sigma_{AKR}^2. \end{aligned} \tag{20}$$

Hence, we get $f_N(y = z - r) \sim N(\mu_N, \sigma_N^2)$ in Eqs. 17 and 18 above. In supplementary Fig. 7 this prediction is shown to be in good agreement with simulations.

For the case of $\mu \neq 0$ we can extract $\mu$ from $A$ as done in Eq. 6 and von Neumann expand the inverse in Eq. 13 to get the fixed point abundances as

$$\mathbf{x}^* \approx (I + \sigma K A)\, K \left( \sigma\mu \sum_{j=1}^{N} x_j \mathbf{1} + \mathbf{r} \right). \tag{21}$$

Where $K$ is a diagonal matrix with $K_i$ on the diagonal, $\mathbf{1}$ a constant vector of ones and $\mathbf{r}$ is a vector of intrinsic growth rates. In this form we see that even if $c = 1$ (making the sum $\sigma\mu\sum_{j=1}^{N} x_j$ constant and equal for every species) nonzero values of $\mu$ in the interaction matrix $A$ will add to the mean of $R_i$ and and thus affect when the first extinction occurs. This will shift the value of $\sigma$ of first extinction to smaller values when $\mu < 0$ and larger values when $\mu > 0$. The effect is increased for $c < 1$ when the sum of abundances is a stochastic variable, and as in the case of $r_i = K_i = 1$, the effect is more pronounced for $\mu > 0$ because of larger species abundances in the sum due to the higher degree of mutualistic interactions in $A$.

### 5 Collapse boundary and persistence

To locate the collapse boundary $\sigma_c$ we use a method connected to persistence – the fraction of viable species for a specific choice of parameters. We estimate persistence by the non-trivial fixed point solutions of $\mathbf{x}^* = (RK - \sigma A)^{-1}\mathbf{r}$, corresponding to specific values of $r_i$, $K_i$ ($r_i$ and $1/K_i$ on the diagonal of the diagonal matrices $R$ and $K$ respectively), and the other GLV parameters. For systems of size $N$ with values of $\sigma > \sigma_f(N)$ – larger standard deviation of interaction strength than for the predicted first extinction boundary – the non-trivial solution will include negative $x_i^*$. Although the positive $x_i^*$ are not the fixed-point abundances of a reduced viable system with biodiversity $n$, the number of positive abundances can be used as an approximation of the persistence. In Fig. 3 in the main text we show the statistics of prediction of the persistence obtained from the non-negative abundances compared to the actual persistence for systems of varying sizes ($N = 20 - 1000$).

The reduced system corresponding to the positive abundances and persistence opens up two ways of predicting collapse. First approach, we use the reduced interaction matrix of positive fixed-point entries $\sigma A^* - R^* K^*$ and calculate collapse as the $\sigma$ where the reduced system loses stability. The second approach is to find the $\sigma$ at which the number of remaining species $n$ (number of positive fixed point entries) would be at the complexity limit introduced by May. This works well since May's limit is known to be a good predictor of collapse when assuming feasibility. For both these approaches, statistics were collected for different realisations in simulations, and we use the average as a predictor of collapse.

#### 6 Estimating initial biodiversity $N_{pred}$

To estimate the size of the initial biodiversity $N_{pred}$ from the measurable quantities of a system in the Extinction Continuum we use a rate of extinction based on the given interaction matrix $A^*$, for the known biodiversity $n$. From $A^*$ we get a the number of non-extinct species for increasing values of $\sigma$ as the positive fixed point solutions to the linear equation

$$\mathbf{x} = \left( \frac{r_i^*}{K_i^*} I^* - \sigma A^* \right)^{-1} \mathbf{r}, \tag{22}$$

thus giving us an average rate of extinction for the community in question. With this rate we extrapolate linearly to the interaction standard deviation of our first extinction prediction for the community $\sigma_f$ (based on the biodiversity of the community $n$), as shown in Fig. 5. This gives a prediction for $N_{pred}$. For systems that are close to collapse the rate obtained in this way tends to diverge. To avoid this we also use an average extinction rate obtained from many runs with systems of all sizes (approximately $rate = -1$), then scaled to fit the specific system size in question $rate\sqrt{cn^3}$. This average rate is used when the predicted specific system rate becomes unrealistically large.

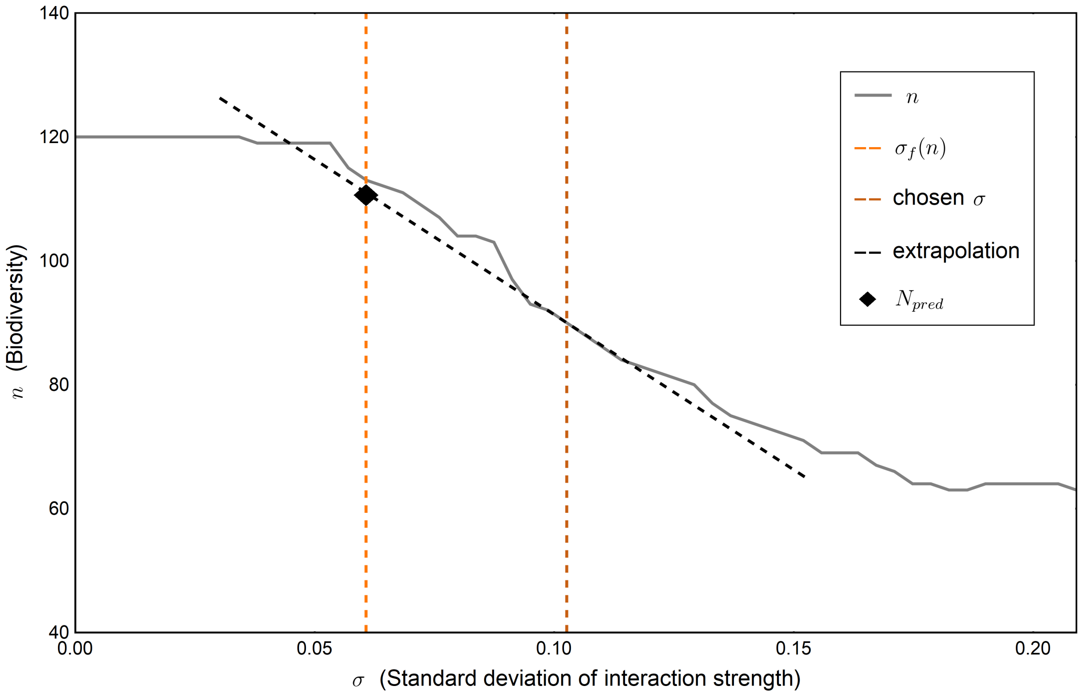


**Figure 5 | Finding $N_{pred}$.** The plot shows the extrapolation (black striped line) with the use of an extinction rate obtained from the interaction matrix $A^*$ for a system in the Extinction Continuum with $n = 90$ species and $\sigma = 0.1$. The size of the initial biodiversity for this system is $N = 120$ and the prediction obtained is $N_{pred} = 111$ marked where the striped orange line indicating the first extinction prediction from $n$ and extrapolation line cross. The grey line is the amount of non-extinct species for the simulated system.

#### 7 The extinction Continuum persists

The Extinction Continuum and associated single species extinction stabilising mechanism hold not only for systems with a standard setup of random interactions between species (no imposed structure, for example, predator-prey), interaction strengths sampled from a normal distribution with a mean of zero and variance of one and of specific sizes as shown in Fig. 2 in the main text. In Figs. 6-10 we show that there are no qualitative differences in results even if any of the assumptions on distribution, structure, mean, $r_i$, and $K_i$ respectively are modified. The figures also show our predictions of the two boundaries surrounding the Extinction Continuum.

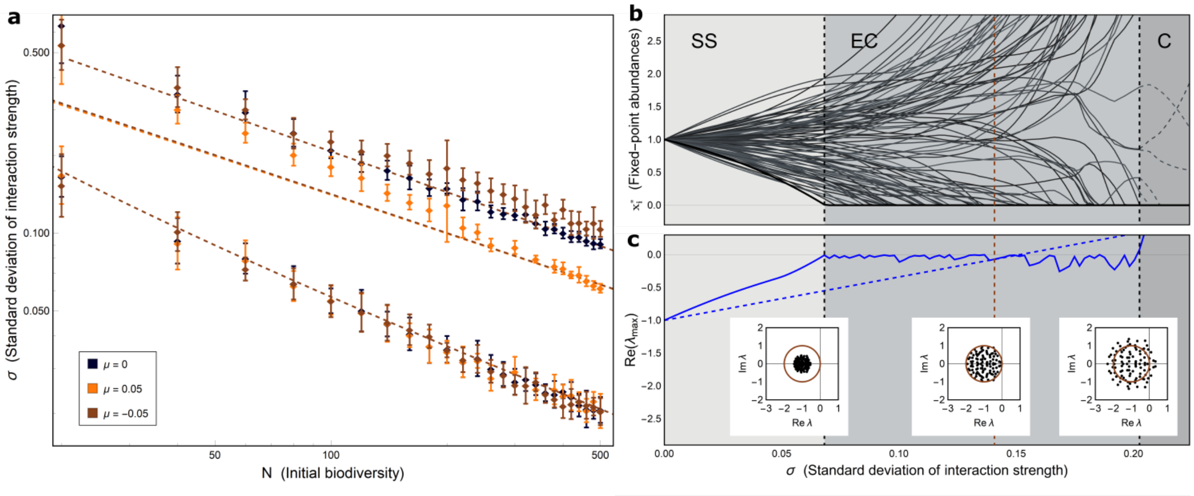


**Figure 6 | Increasing interaction variability in complex systems with $\mu \neq 0$.** (**a**) Is a log-log plot showing standard deviation of interaction strength for first extinction, extended May's limit [5] and collapse vs. initial biodiversity $N$ for a random system with Normally distributed interaction strengths with $\mu = 0$ (black) and two systems with $\mu = -0.05$ (brown) and $\mu = 0.05$ (yellow) respectively. The dots are simulated values with one standard deviation errorbars and the striped lines are our theoretical predictions. All systems have intrinsic growth rates and carrying capacities $r_i = k_i = 1$ and connectance $c = 0.5$. (**b,c**) Show a specific run for a system with $\mu = -0.05$, $c = 0.5$ and $N = 100$. (**b**) Shows fixed-point species abundances $x_i^*$ for varying $\sigma$. The shaded background indicates the three phases Strict Stability (SS), Extinction Continuum (EC) and Collapse (C). The three dashed lines show the two boundaries first extinction and collapse with the limit introduced by May in between. (**c**) Shows the the leading eigenvalue of the system (blue line) and the leading eigenvalue for the same system when assuming feasibility (not including fixed point species abundances in the Jacobian), blue dotted line. The insets show the spectrum of the Jacobian excluding the species abundances at the two boundaries and May's limit with the circle indicating the stability radius. It is clearly seen that the Extinction Continuum with single species extinctions to uphold community resilience is present also in the case for $\mu \neq 0$.

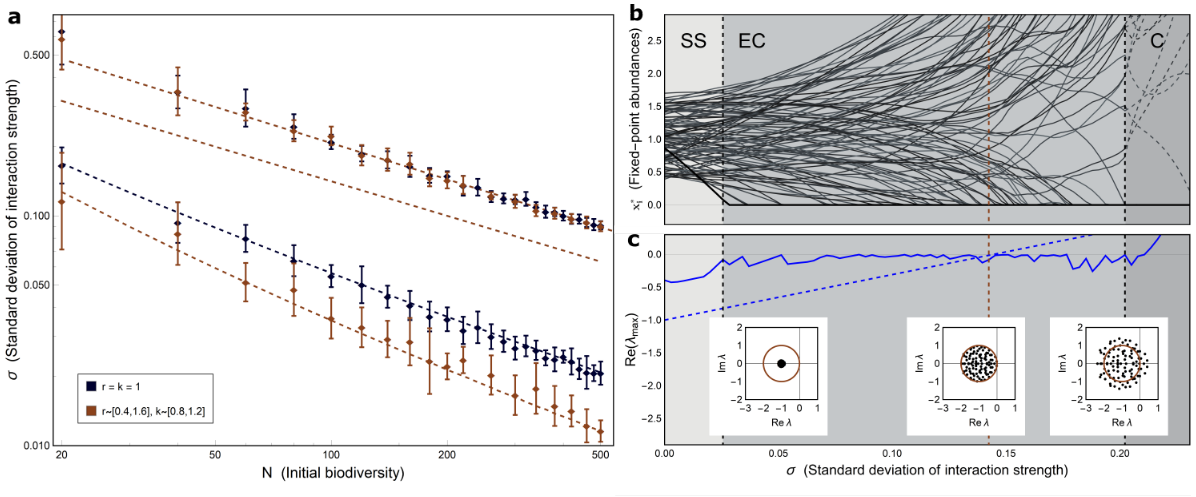


**Figure 7 | Increasing interaction variability in complex a system for $r_i$ and $k_i$ with non-zero variance.** (**a**) Is a log-log plot showing standard deviation of interaction strength for first extinction, May's limit and collapse vs. initial biodiversity $N$ for a random system with $r_i = k_i = 1$ (black) and a system with $r_i$ and $k_i$ drawn from uniform distributions Unif$(0.4, 1.6)$ and Unif$(0.8, 1.2)$ respectively (brown). The dots are simulated values with one standard deviation errorbars and the striped lines are our theoretical predictions. Both systems have Normal$(0, 1)$ for the entries in the interaction matrix and connectance $c = 0.5$. (**b,c**) Show a specific run for a system with varying $r_i$ and $k_i$, $c = 0.5$ and $N = 100$. (**b**) Shows fixed point species abundances $x_i^*$ for varying $\sigma$. The shaded background indicates the three phases Strict Stability (SS), Extinction Continuum (EC) and Collapse (C). The three dashed lines show the two boundaries first extinction and collapse with the limit introduced by May in between. (**c**) Shows the the leading eigenvalue of the system (blue line) and the leading eigenvalue for the same system when assuming feasibility (not including fixed point species abundances in the Jacobian), blue dotted line. The insets show the spectrum of the Jacobian excluding the species abundances at the two boundaries and May's limit with the circle indicating the stability radius. It is clearly seen that the Extinction Continuum is present even in cases when $r_i$ and $k_i$ are drawn from distributions. Although with a large enough variance in $r_i$ the SBS phase will disappear.

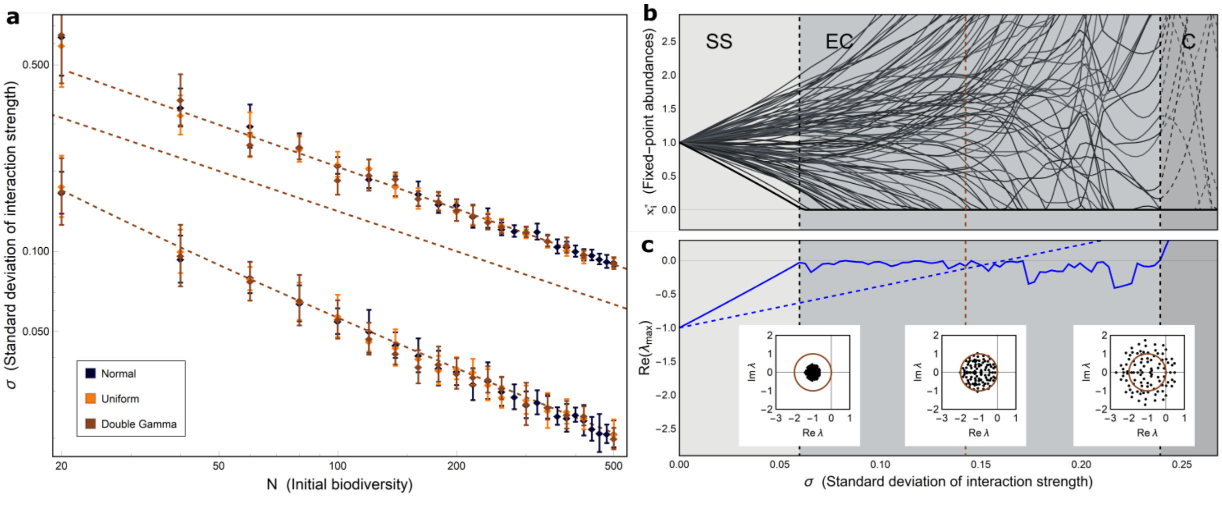


**Figure 8 | Increasing interaction variability in complex systems with different interaction strength distributions.** (**a**) Is a log-log plot showing standard deviation of interaction strength for first extinction, May's limit and collapse vs. initial biodiversity $N$ for a random system with interaction strength distribution Normal$(0, 1)$ (black) and two systems with Uniform$(-1/\sqrt{3}, 1/\sqrt{3})$ (yellow) and a mirrored Gamma distribution MirrorGamma$(0, 1)$ respectively. The dots are simulated values with one standard deviation errorbars and the striped lines are our theoretical predictions. All systems have intrinsic growth rates and carrying capacities $r_i = k_i = 1$ and connectance $c = 0.5$. (**b,c**) Show a specific run for a system with MirrorGamma$(0, 1)$ interaction strength distribution, $c = 0.5$ and $N = 100$. (**b**) Shows fixed point species abundances $x_i^*$ for varying $\sigma$. The shaded background indicates the three phases Strict Stability (SS), Extinction Continuum (EC) and Collapse (C). The three dashed lines show the two boundaries first extinction and collapse with the limit introduced by May in between. (**c**) Shows the the leading eigenvalue of the system (blue line) and the leading eigenvalue for the same system when assuming feasibility (not including fixed point species abundances in the Jacobian), blue dotted line. The insets show the spectrum of the Jacobian excluding the species abundances at the two boundaries and May's limit with the circle indicating the stability radius. It is clearly seen that Extinction Continuum is present also for different interaction strength distributions. It has been previously shown that as long as the distributions have the same mean and variance the will have equal May limit **??**, our boundary predictions show that this also holds for first extinction and the collapse boundary.

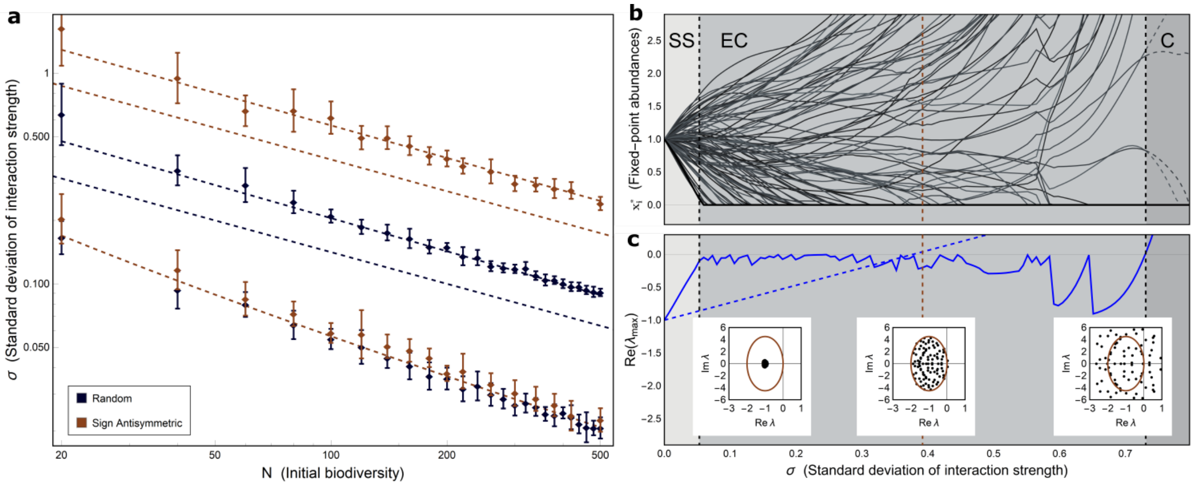


**Figure 9 | Increasing interaction variability in a complex system with predator-prey structure.** (**a**) Is a log-log plot showing standard deviation of interaction strength for first extinction, extended May's limit [5] and collapse vs. initial bidiversity $N$ for a random system without any structure in the interaction matrix (black) and a system with predator-prey structure sign($a_{ij}$)=-sign($a_{ji}$) (brown). The dots are simulated values with one standard deviation errorbars and the striped lines are our theoretical predictions. All systems have intrinsic growth rates and carrying capacities $r_i = k_i = 1$, Normally distributed interaction strengths Normal$(0, 1)$ and connectance $c = 0.5$. (**b,c**) Show a specific run for a system with predator-prey structure and $N = 100$. (**b**) Shows fixed point species abundances $x_i^*$ for varying interaction variability. The shaded background indicates the three phases Strict Stability (SS), Extinction Continuum (EC) and Collapse (C). The three dashed lines show the two boundaries first extinction and collapse with the limit introduced by May in between. (**c**) Shows the the leading eigenvalue of the system (blue line) and the leading eigenvalue for the same system when assuming feasibility (not including fixed point species abundances in the Jacobian), blue dotted line. The insets show the spectrum of the Jacobian excluding the species abundances at the two boundaries and May's limit with the oval indicating the stability radius. It is clearly seen that including persistence in the analysis radically changes stability predictions introducing the Extinction Continuum also for systems with predator-prey structure.

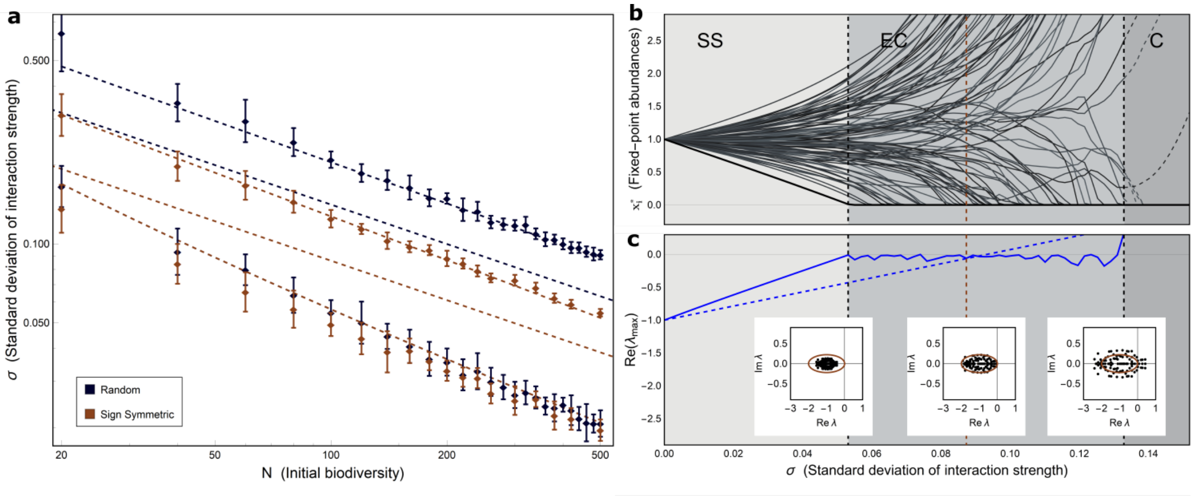


**Figure 10 | Increasing interaction variability in a complex system with mutualistic and competitive interactions.** (**a**) Is a log-log plot showing standard deviation of interaction strength for first extinction, extended May's limit [5] and collapse vs. initial biodiversity $N$ for a random system without any structure in the interaction matrix (black) and a system with only mutualistic and competitive interactions sign($a_{ij}$)=sign($a_{ji}$) (brown). The dots are simulated values with one standard deviation errorbars and the striped lines are our theoretical predictions. All systems have intrinsic growth rates and carrying capacities $r_i = k_i = 1$, Normally distributed interaction strengths Normal$(0, 1)$ and connectance $c = 0.5$. (**b,c**) Show a specific run for a system with predator-prey structure and $N = 100$. (**b**) Shows fixed-point species abundances $x_i^*$ for varying $\sigma$. The shaded background indicates the three phases Strict Stability (SS), Extinction Continuum (EC) and Collapse (C). The three dashed lines show the two boundaries first extinction and collapse with the limit introduced by May in between. (**c**) Shows the the leading eigenvalue of the system (blue line) and the leading eigenvalue for the same system when assuming feasibility (not including fixed point species abundances in the Jacobian), blue dotted line. The insets show the spectrum of the Jacobian excluding the species abundances at the two boundaries and May's limit with the oval indicating the stability radius. It is clearly seen that including persistence in the analysis changes the stability predictions and introduces the Extinction Continuum also for cases when species interactions are strictly mutualistic or competitive.

### 8 The case of large distribution mean $|\mu|$

As noted in the main text section 2.2 the $\gamma$ measure of proximity to collapse is not reliable for a mean of the interaction strengths with significantly larger absolute value than zero. This is because we estimate $\sigma_c$ (loss of stability) based on when the matrix ($\sigma A^* - D^*$, with $D^*$ a diagonal matrix with $r_i/K_i$ on the diagonal) for a reduced system estimated by the positive entries of the non-trivial solution to equation

$$\mathbf{x} = \left( \frac{r_i^*}{K_i^*} I^* - \sigma A^* \right)^{-1} \mathbf{r}^*, \tag{23}$$

loses stability. This is not the actual Jacobian of the reduced system which would be $X^*(\sigma A^* - D^*)$, where $X^*$ is a diagonal matrix with the fixed point abundances of the reduced system on the diagonal. Not including the fixed-point abundances (since they are feasible for the reduced systems and assumed not to alter stability outcome with D-stability arguments [**RN147**, 6]) is the approach that May and many after him use to approximate the collapse boundary. This is a more general approach since it does not need the species abundances and therefore not the specification of a system.

Generality on the other hand has its costs, even in addition to not including extinctions. In our boundary predictions we are taking extinctions into account by using the persistence, but there is a difference in magnitude of the fixed-point abundances of the real (simulated) system depending on sign and magnitude of a non-zero $\mu$. These magnitude differences in species abundances acts either to destabilise or stabilise the Jacobian which is not accounted for since we approximate persistence without abundances. For systems with $\mu > 0$ there is a prevalence of mutualistic interactions making the fixed point abundances larger and thus destabilising the Jacobian for smaller values of $\sigma$. Equally when $\mu < 0$ there will be more predator-prey interactions keeping the abundances in check which stabilises the Jacobian leading to a collapse boundary at larger $\sigma$. It is an interesting observation that simulated systems with larger non-zero $|\mu|$ do not, as for the $\mu \approx 0$ case, agree with the theoretical predictions for larger $|\mu|$ [5]. For these reduced systems collapse occurs at larger/smaller values of $\sigma$ due to the inclusion of species abundances.
